# Insulin-mediated suppression of fatty acid release predicts whole-body insulin resistance of glucose uptake and skeletal muscle insulin receptor activation

**DOI:** 10.1101/2024.02.29.582589

**Authors:** Michael W Schleh, Benjamin J Ryan, Cheehoon Ahn, Alison C Ludzki, Douglas W Van Pelt, Lisa M Pitchford, Olivia K Chugh, Austin T Luker, Kathryn E Luker, Dmitri Samovski, Nada A Abumrad, Charles F Burant, Jeffrey F Horowitz

## Abstract

To examine factors underlying why most, but not all adults with obesity exhibit impaired insulin-mediated glucose uptake, we compared: 1) rates of fatty acid (FA) release from adipose tissue, 2) skeletal muscle lipid droplet (LD) characteristics, and 3) insulin signaling events in skeletal muscle collected from cohorts of adults with obesity with “HIGH” versus “LOW” insulin sensitivity for glucose uptake. Seventeen adults with obesity (BMI: 36±3kg/m^2^) completed a 2h hyperinsulinemic-euglycemic clamp with stable isotope tracer infusions to measure glucose rate of disappearance (glucose Rd) and FA rate of appearance (FA Ra). Skeletal muscle biopsies were collected at baseline and 30min into the insulin infusion. Participants were stratified into HIGH (n=7) and LOW (n=10) insulin sensitivity cohorts by their glucose Rd during the hyperinsulinemic clamp (LOW<400; HIGH>550 nmol/kgFFM/min/[µU/mL]). Insulin-mediated suppression of FA Ra was lower in LOW compared with HIGH (p<0.01). In skeletal muscle, total intramyocellular lipid content did not differ between cohorts. However, the size of LDs in the subsarcolemmal region (SS) of type II muscle fibers was larger in LOW compared with HIGH (p=0.01). Additionally, insulin receptor (IR) interactions with regulatory proteins CD36 and Fyn were lower in LOW versus HIGH (p<0.01), which aligned with attenuated insulin-mediated Tyr phosphorylation of IRβ and downstream insulin-signaling proteins in LOW. Collectively, reduced ability for insulin to suppress FA mobilization, with accompanying modifications in intramyocellular LD size and distribution, and diminished IR interaction with key regulatory proteins may be key contributors to impaired insulin-mediated glucose uptake commonly found in adults with obesity.

**KEY POINTS:**

- Although most adults with obesity exhibit impaired insulin-mediated glucose uptake (insulin resistance), some remain sensitive to insulin. Factors that “protect” adults with obesity from developing resistance to insulin-mediated glucose uptake are poorly understood.
- Potent suppression of fatty acid (FA) mobilization from adipose tissue by insulin is a strong predictor of whole-body insulin-mediated glucose uptake.
- Participants with higher sensitivity for insulin-mediated glucose uptake had smaller intramyocellular lipid droplets (LDs) within the subsarcolemmal region of type II skeletal muscle fibers.
- Novel findings revealed that insulin receptor (IR) interaction with the long-chain fatty acid transport protein, CD36, and the Src-family kinase, Fyn, directly associated with higher rates of glucose uptake under basal and hyperinsulinemic conditions.
- Together, the findings suggest impaired suppression of FA release from adipose tissue associates with reduced glucose uptake in skeletal muscle due in part to a defect in IR activation by CD36/Fyn and altered subcellular LD characteristics.

## INTRODUCTION

Obesity has now surpassed 40% of the United States population (1), and is a major risk factor for the development of chronic metabolic conditions such as insulin resistance (2), which in turn underlies many obesity-related diseases (3). Although most adults with obesity are insulin resistant for glucose uptake, some remain relatively insulin sensitive (4, 5), and factors helping to protect against the development of insulin resistance in these individuals are not clear. Work from our lab (6, 7) and others (8-10) demonstrated the rate of fatty acid (FA) mobilization into the systemic circulation is directly linked with whole-body insulin sensitivity among adults with obesity. The vast majority of FA in the systemic circulation are derived from abdominal subcutaneous adipose tissue (aSAT) (11), and we contend the variability in the regulation of FA release from aSAT is an important contributor to the differences in the magnitude of insulin resistance observed among adults with obesity.

The link between excessive systemic FA mobilization and whole-body insulin resistance has been largely attributed to high rates of FA uptake into the intramyocellular region of skeletal muscle and the resultant over-accumulation of bioactive lipids (12-15). In particular, some bioactive lipids reported to promote insulin resistance include diacylglycerol (DAG) localized to the muscle membrane (16), total ceramide (17) and long-chain fatty acyl-CoA (18, 19). The over-accumulation of these bioactive lipids are proposed to activate PKC family proteins (PKC ε, θ, ζ), resulting in Ser/Thr phosphorylation of insulin receptor substrate-1 (IRS-1), and reduced phosphorylation of AKT (20-22). However, the contribution of each of these lipid intermediates to insulin resistance has been challenged (23-25), and PKC-independent mechanisms may also underlie poor activation in the development of insulin resistance. Intramyocellular lipids are primarily stored within lipid droplets (LDs) composed of a triacylglycerol-rich core surrounded by a phospholipid monolayer containing proteins that regulate FA kinetics and direct intracellular LD trafficking (26, 27). Importantly, the metabolic impact of LDs can vary depending on their location within the myocyte. For example, LDs within the intramyofibrillar region (IMF; towards the center of the myocyte) are proposed to support mitochondrial respiration to help meet energy requirements during muscle contraction, and LDs in the subsarcolemmal region (SS; towards the periphery of the myocyte) primarily support the metabolic demand from the plasma membrane (28, 29). The intracellular distribution of LDs within the myocyte has also been linked with insulin resistance, where LD accumulation in the SS region has been associated with insulin resistance and type 2 diabetes (30-32), perhaps a consequence of a greater lipid intermediate release in the vicinity of the membrane where insulin signaling is initiated. Differences in the number and size of intramyocellular LDs within the IMF and SS regions may also impact insulin sensitivity, but these relationships are not firmly established. One of the principal aims of this study was to compare the number and size of LDs within the IMF and SS regions of type I and type II skeletal muscle fibers from a cohort of adults with HIGH versus LOW insulin sensitivity.

The disruption of insulin signaling induced by the intramyocellular lipids have classically been attributed to modifications in the phosphorylation of signaling proteins downstream of the insulin receptor (IR) (e.g., IRS-1, phosphoinositide 3-kinase (PI3K), and AKT). However, other reports suggest lipid-induced disruption directly at the IR may also contribute to impaired glucose uptake (33). The membrane glycoprotein, cluster of differentiation 36 (CD36) is well-known for its role in regulating long-chain FA transport (34-36), but has also been implicated in functions related to cellular metabolism and IR phosphorylation (33, 37), thereby modifying downstream signaling (38). The physical interaction between CD36 and the IR has been found to increase Tyr phosphorylation of the IR via CD36-mediated recruitment of the Src-family tyrosine kinase, Fyn (33). Conversely, previous reports demonstrated elevated saturated FA availability attenuated Fyn recruitment to the CD36-IR complex (33, 38, 39), leading to blunted insulin signaling (33, 38). These previous experiments were conducted *in vitro*, and whether differences in systemic FA availability among adults with obesity may contribute to modifications in CD36 recruitment of Fyn to the IR and modify IR phosphorylation and downstream insulin action in human skeletal muscle is not known. The second major aim of this study was to compare basal- and insulin-mediated interaction between CD36 and Fyn with the IR, as well as insulin-stimulated signaling events in skeletal muscle from a cohort of adults with high versus low insulin sensitivity.

## METHODS

### Participants

Seventeen men (n=6) and women (n=11) with obesity participated in this study (BMI=30-40 kg/m^2^). All participants were ‘inactive’ by not engaging in any planned physical aerobic or resistance exercise, and weight stable (±2 kg for the previous 6 months). Participants did not take medications known to affect glucose or lipid metabolism, have a history of heart disease, or actively smoke. All women participating in the study were pre-menopausal with regularly occurring menses, and not pregnant or lactating. Participants provided written informed consent before participation. The study conformed to the standards set by the *Declaration of Helsinki*. The study protocol was approved by the University of Michigan Institutional Review Board and was registered at clinicaltrials.gov (NCT02717832). Participants in the present study also participated in a previous study from our laboratory focused on metabolic dysfunction in adipose tissue (7). Findings presented here are independent from our previously published work.

### Preliminary assessment: body composition, visceral fat area, and liver fat

Body composition was assessed by dual-energy X-ray absorptiometry (Lunar DPX DEXA Scanner, GE, Madison, WI) at the Michigan Clinical Unit (MCRU). Visceral fat area (cm^2^) and hepatic fat percentage was measured by magnetic resonance imaging (MRI: Ingenia 3T MR System, Phillips, Netherlands) at the University of Michigan Medicine’s Department of Radiology.

### Experimental protocol

The night before the experimental trial, participants were provided a standardized meal at 1900h and a snack at 2200h, containing 30% and 10% of their estimated daily energy expenditure, respectively (Figure 1). The macronutrient composition of both the meal and snack were 55% carbohydrate, 30% fat, 15% protein. After the evening snack the participants remained fasted until completion of the study trial the next day. The next morning, participants arrived to MCRU at 0700h and quietly rested for 60min. Intravenous catheters were inserted at ∼0800h into a hand vein for continuous blood sampling, and forearm vein for continuous isotope, insulin, and glucose infusion. At ∼0900h, a baseline blood sample was collected for background isotope labeling, followed by a primed continuous infusion of [6,6 ^2^H_2_]glucose (35 µmol/kg priming dose, and 0.41 µmol/kg/min continuous infusion; Cambridge Isotopes, Tewksbury, MA). At ∼0915h, a skeletal muscle biopsy was collected from the vastus lateralis muscle, and visible connective tissue and lipid were quickly removed. A portion of each muscle sample was preserved for histology by manually aligning the muscle fibers in parallel under magnification before embedding in optimal cutting temperature (OCT) compound and then flash freezing in isopentane chilled over liquid nitrogen. The remaining portion of the muscle samples were flash frozen in liquid nitrogen for immunoblot analysis. After the basal muscle biopsy, a continuous [1-^13^C]palmitate bound to human albumin infusion began at ∼1000h (0.04 µmol/kg/min continuous infusion; Cambridge). By ∼1050h, three ‘arterialized’ samples were collected (heated-hand technique) for measurements of basal glucose and FA kinetics. At ∼1100h, a hyperinsulinemic-euglycemic clamp procedure began (40 mU insulin/m^2^/min) for determination of whole-body insulin sensitivity (40). Blood samples were collected every 5min during the clamp to measure blood glucose concentration (StatStrip, Nova Biomedical, Waltham MA) and the infusion rate of dextrose (20% dextrose in 0.9% NaCl) was adjusted accordingly to accommodate insulin-induced hypoglycemia and attain baseline glucose concertation (∼5.0 mM). Exactly 30min into the hyperinsulinemic-euglycemic clamp procedure, a second muscle biopsy was obtained to identify insulin-mediated signaling events occurring after acute insulin exposure. After plasma glucose concentration stabilized for a minimum of 20min without adjusting glucose infusion rate (∼120min into the clamp), arterialized blood samples were collected to quantify insulin-mediated glucose and FA kinetics.

**Figure 1:**
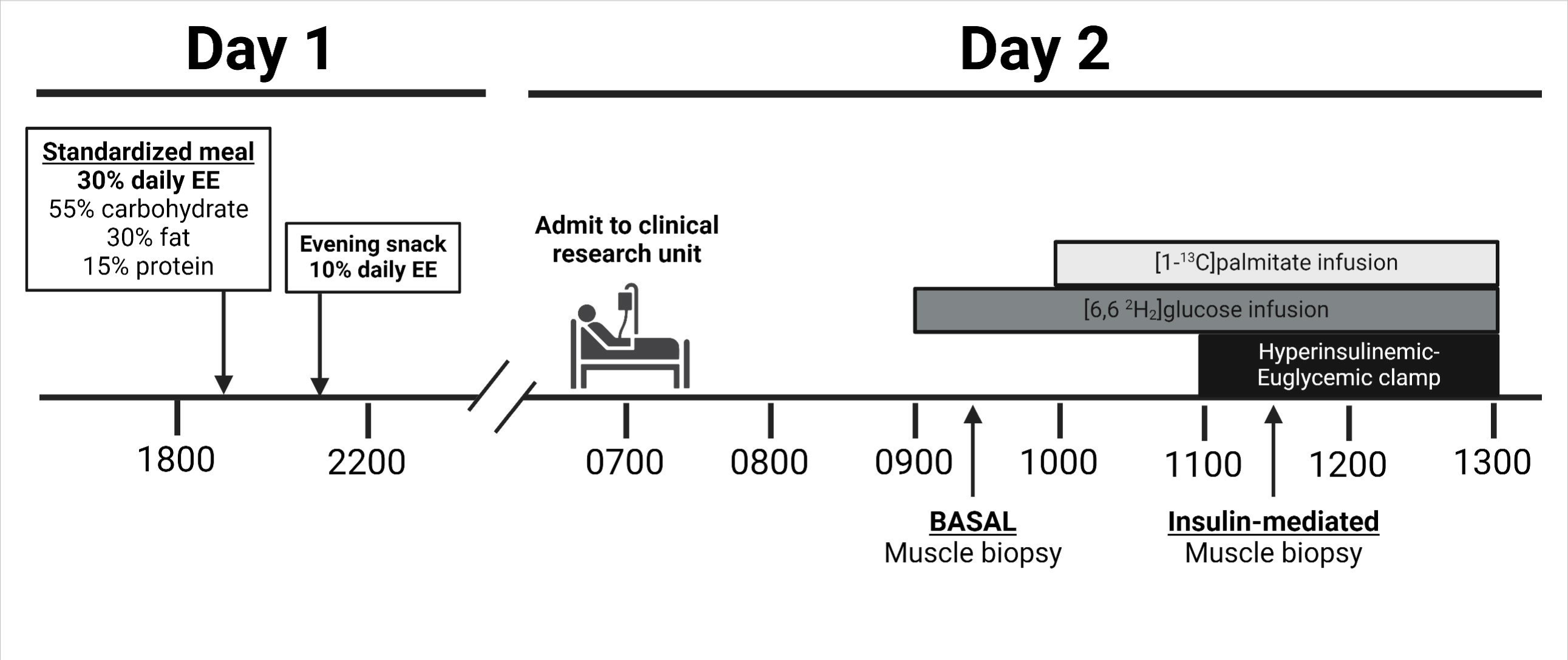
Clinical trial timeline of sample collection. Insulin-mediated skeletal muscle biopsies were collected 30min into the hyperinsulinemic clamp. EE, energy expenditure. Created with BioRender.com

### Participant stratification by glucose rate of dissapearance

For paired analyses, participants were stratified into two cohorts: LOW glucose rate of disappearance (Rd) relative to circulating steady-state insulin at the end of the clamp (Rd/I: 181 nmol/kg FFM/min / (µU/mL) ≤ LOW ≤ 382 nmol/kg FFM/min / (µU/mL); n=10) and HIGH glucose Rd/I (552 nmol/kg FFM/min / (µU/mL) ≤ HIGH ≤ 1274 nmol/kg FFM/min / (µU/mL); n=7). To confirm our measures and participant stratification, HOMA-IR for all members of the LOW cohort was > 2.5, which is a recognized index of peripheral insulin resistance [Table 1; (41)].

**Table 1:** Participant characteristics.

|  | **HIGH (n=7)** | **LOW (n=10)** | **P-value** |
| --- | --- | --- | --- |
| **Sex (M/F)** | 1/6 | 5/5 |  |
| **Age (yrs)** | 32 ± 9 | 31 ± 8 | 0.75 |
| **Body mass (kg)** | 93 ± 9 | 105 ± 11* | 0.02 |
| **BMI (kg/m<sup>2</sup>)** | 35 ± 3 | 36 ± 3 | 0.18 |
| **Fat mass (kg)** | 43 ± 5 | 46 ± 5 | 0.22 |
| **Fat free mass (kg)** | 50 ± 10 | 60 ± 9 | 0.06 |
| **Body fat %** | 46 ± 6 | 43 ± 5 | 0.22 |
| **Liver fat (%)** | 3.4±2.0 | 9.5±8.3 | 0.19 |
| **Visceral fat (cm<sup>2</sup>)** | 107±49 | 169±58 | 0.10 |
| **Plasma Parameters** |  |  |  |
| **Fasting plasma glucose (mM)** | 4.6 ± 0.4 | 4.7 ± 0.4 | 0.71 |
| **Fasting plasma insulin (μU/mL)** | 6.3 ± 2.5 | 16.8 ± 4.7* | 3.4e<sup>-05</sup> |
| **HOMA-IR** | 1.3 ± 0.6 | 3.5 ± 1.0* | 6.3e<sup>-05</sup> |
| **HbA1c (%)** | 5.3 ± 0.4 | 5.4 ± 0.2 | 0.73 |
| **Plasma FA (mM)** | 438 ± 197 | 544 ± 176 | 0.27 |
| **Plasma triacylglycerol (mg/dL)** | 58 ± 34 | 80 ± 38 | 0.24 |
| **Plasma HDL (mg/dL)** | 44 ± 10 | 38 ± 12 | 0.27 |
| **Plasma cholesterol (mg/dL)** | 141 ± 15 | 143 ± 32 | 0.88 |
| **Plasma CRP (mg/L)** | 11 ± 14 | 9 ± 11 | 0.79 |
| **Plasma IL6 (pg/mL)** | 3.0 ± 1.4 | 3.3 ± 2.2 | 0.74 |
| **Plasma HMW adiponectin (ng/mL)** | 2581 ± 1412 | 1658 ± 1261 | 0.09 |
| **Plasma adiponectin (Total) (ng/mL)** | 5978 ± 2139 | 3840 ± 2017 | 0.06 |
| **Plasma leptin (ng/mL)** | 61 ± 29 | 51 ± 19 | 0.44 |

Data are mean ± SD. BMI, body mass index. CRP, C-reactive protein. FA, fatty acid. HDL, high density lipoprotein. HIGH, high insulin sensitivity. HMW adiponectin, high molecular weight adiponectin. HOMA-IR, homeostatic model for insulin resistance. LOW, low insulin sensitivity \*p<0.05 vs HIGH.

## ANALYTICAL PROCEDURES

### Hepatic and visceral fat

Liver and visceral fat content were measured from MRI images by chemical-shift-encoded MRI proton density fat fraction (42), and acquisition as previously described by our lab (7, 43). Visceral fat area was measured from 3 separate axial sections (5mm thickness) in the L2-L3 vertebral region, and hepatic fat percentage was measured from 3 separate axials slices (5mm thickness) from the liver. Both liver and visceral fat area were quantified by a trained investigator blinded to participant identification.

### Plasma glucose and fatty acid kinetics

The tracer-to-tracee ratio (TTR) for calculations pertaining to glucose and palmitate kinetics were determined by gas-chromatography-mass spectrometry (GC/MS) using a Mass Selective Detector 5973 system (Agilent Technologies, Santa Clara, CA), as described by our lab (7, 44). FA rate of appearance (Ra) was quantified by dividing palmitate Ra by the ratio of total palmitate (C16:0) to FAs within a standard. Glucose TTR was selectively analyzed and peak abundances were averaged at the following m/z for ion fragments (103/105, 115/117, 127/129, and 217/219) for endogenous and isotope labeled [6,6 ^2^H_2_]glucose respectively. FA Ra and glucose Ra were calculated from their respective TTR using the Steele equation for steady state conditions (45). Because samples were collected during steady-state conditions, rates of disappearance from the circulation were equal to their rates of appearance into the circulation (i.e., Ra = Rd).

### Plasma glucose, lipid, and insulin concentration

Plasma glucose, non-esterified fatty acid, triacylglycerol, high-density lipoprotein (HDL), and total cholesterol were measured in plasma from commercially available colorimetric assays. Plasma interleukin-6 (IL-6), C-reactive protein (CRP), leptin, total adiponectin, and high molecular-weight (HMW) adiponectin were measured via enzyme-linked immunoassay (ELISA). Insulin was measured in serum by radioimmunoassay (Siemens). See Table S1 for vendor details.

### Skeletal muscle histochemistry: LD processing and fluorescence image acquisition and analysis

Muscle samples from OCT embedded tissues were cut into 5µm sections at - 20°C and mounted onto an ethanol-cleaned glass slide. Sections were fixed in 4% paraformaldehyde for 60min, and permeabilized for 5min in 0.5% triton X-100. Sections were then incubated with primary antibodies myosin heavy chain type 1 (MHC-1) at 1:200 for 60min at 37°C, followed by 30min secondary antibody incubation with AlexaFluor 647, goat-anti mouse IgG2b at 1:200. Afterwards, sections were stained with Bodipy 493/503 at 1:100 for 20min, followed by 20min incubation with anti-wheat germ agglutinin (WGA) conjugated with AlexaFluor 555 at 1:500 to identify muscle membranes. Sections were mounted with ProLong Gold Antifade Mountant, covered with #1 coverslips (2975-245, Corning, Corning, NY), and stored in the dark until imaging. See Table S1 for detailed histochemistry vendor details.

Images used for the quantification of LD area (% stained), LD number (number of LDs/mm^2^), and median LD size (µm^2^) were captured on a Keyence Bz-X710 fluorescence microscope (Keyence; Osaka Japan) with a x20 N.A.=0.45 objective lens using 1440 × 1080 pixels (533 µm x 400 µm) field of view. Following image acquisition, LD characteristics (i.e., number and size), and distribution were quantified using MATLAB R2021a (Mathworks, Inc., Natick, MA). In brief, images captured at x20 were converted to grayscale and re-scaled to accommodate image variability between samples. Individual muscle fibers were identified and labeled using a custom ridge detection algorithm and watershed segmentation to complete non-continuous cell borders and completely identify cells. Type I fibers were identified based on positive MHC type I-positive stain, and non-stained fibers considered type II fibers. Fiber types were partitioned into type I or type II fibers dependent on signal intensity using k-means clustering (k=2). Lipid droplets identified in the SS region were contained within ∼3 µm of the peripheral border of each fiber, while the IMF region remained towards the center of each fiber (∼91% of total myocyte volume). Lipid droplets were detected as the mean area of puncta, and a top-hat filter was used to accommodate for image variability and background noise produced by light scatter of the fluorescence microscope. Lipid droplets identified and included for analysis within each muscle fiber were contained within a normal Gaussian distribution. In total, 3-5 fields of view were obtained per participant, resulting in the analysis of 125±57 muscle fibers per participant.

### Skeletal muscle lysate preparation for immunoblot

Frozen muscle samples were weighed (∼25mg) and transferred into pre-chilled microcentrifuge tubes containing 1mL ice-cold lysis buffer (RIPA: 20 mM Tris-HCl-pH 7.5, 150 mM NaCl, 1 mM Na_2_EDTA, 1 mM EGTA, 1% NP-40, 1% sodium deoxycholate, 2.5 mM sodium pyrophosphate, 1 mM β-glycerophosphate, 1 mM Na_3_VO_4_, 1 µg/mL leupeptin [9086, Cell Signaling Technology, Danvers, MA]), 1% phosphatase inhibitor cocktail #1 & #2 (P2850, P5726, Sigma-Aldrich, St. Louis, MS), 1% protease inhibitor cocktail (P8340, Sigma-Aldrich), and two steel ball bearings per sample. Samples were homogenized for 30s at 45Hz using Qiagen TissueLyser II, then solubilized for 60min by inverted end-over-end rotation at 50rpm in the cold (4°C). Afterwards, samples were centrifuged at 15000g for 15min in 4°C, discarding the pellet. Protein concentration was determined by bicinchoninic acid (BCA) assay (23225, Thermo-Fisher, Waltham, MA), and samples were diluted in 4x Laemmli buffer (1610747, Bio-Rad, Hercules, CA) to 1mg/mL concentration, then heated for 7min in 95°C. Proteins were separated by SDS-Page (8-12% acrylamide), transferred to nitrocellulose membranes, and probed for the primary antibodies described in Table S1. To account for loading control, membranes were normalized to total protein abundance determined by Memcode reversible protein stain (Pierce Thermo-Fisher). Additionally, each gel contained an internal standard sample from 8 obese individuals to account for gel-to-gel variability, in which each sample was also normalized to their internal standard.

### Immunoprecipitation

Frozen skeletal muscle samples were homogenized in 1% triton-based lysis buffer (20 mM Tris-HCl (pH 7.5), 150 mM NaCl, 1 mM Na2EDTA, 1 mM EGTA, 1% triton X-100, 2.5 mM sodium pyrophosphate, 1 mM beta-glycerophosphate, 1 mM Na3VO4, 1 µg/ml leupeptin); 9803, Cell Signaling Technology) using a chilled dounce homogenizer. Homogenates were then solubilized by end-over-end rotation for 60min, and centrifuged at 100000g for 60min to limit sarcomere proteins. The supernatant was then collected, protein concentration quantified (BCA), and 300µg protein from the lysate was adjusted to 500µL volume with lysis buffer. The adjusted volume was rotated end-over-end overnight at 4°C with 5µg IRβ antibody. The following day, 25µL protein-A magnetic beads (88845, Pierce Thermo-Fisher) were washed in wash buffer (25 mM Tris, 0.15M NaCl, 0.05% Tween-20, pH 7.5), then added to the lysate-antibody mixture, and rotated gently end-over-end for 120min at room temperature. The immunoprecipitation matrix was separated using a magnetic rack (DynaMag2, 12321D, Thermo-Fisher) and washed three times with lysis buffer, twice with wash buffer, and once with PBS. Antigens were eluted from the bead complex with 80 µL 2X Laemmli SDS buffer, by a combination of rotation and vortexing samples for 10min, and discarding the beads following protein elution. Following immunoprecipitation, protein abundance from the eluates was analyzed by immunoblot for basal and insulin-mediated CD-36, IRβ, and Fyn. CD36 and Fyn interaction with IRβ was normalized to a 15µL input sample from the same participant loaded onto the same gel as the eluate.

### Calculations

#### Insulin-mediated glucose uptake

Insulin-mediated glucose uptake was quantified by rate of glucose Rd at the end of the hyperinsulinemic-euglycemic clamp. Because glucose was in “steady-state” during the final 20min of the clamp, glucose Rd at this time was calculated as the sum of the exogenous glucose infusion rate and endogenous hepatic glucose production (glucose Ra). Glucose Rd was then normalized to both fat-free mass (FFM) and the steady-state insulin concentration (endogenous insulin production + exogenous insulin infusion) at the end of the clamp (Rd/I = nmol glucose/kg FFM/min / [µU/mL insulin]).

#### Statistical analysis

All measurements that were not normally distributed were log-transformed. Linear mixed models were used to examine the main effects for group (HIGH versus LOW), main effects for insulin-stimulation (basal versus insulin-mediated conditions), and group x insulin interaction for measurements including targeted protein analysis, immunoprecipitation, FA Ra, and glucose Ra. For LD histochemistry analysis, measurements of lipid content, LD number, and LD area were segregated into regional distribution (IMF and SS region). Linear mixed models were then used on the segregated outcomes to identify main effects of group (HIGH versus LOW), main effects for fiber type (type I vs type II), and group x fiber-type interaction in the IMF and SS region independently. When significant effects were observed, Fisher’s least significant difference post-hoc test was used to identify significant interactions between groups. Unpaired Student’s t-tests were used to test for significant differences between groups (HIGH versus LOW) for all other clinical variables. Pearson’s correlation was used to identify relationships between clinical outcomes and glucose Rd/I. Statistical analysis was completed using R version 4.1.0. Data are displayed as mean ± SD, and significance set to p<0.05.

## RESULTS

### Cohort stratification based on insulin-mediated glucose uptake

As anticipated, insulin-mediated glucose uptake (measured as glucose Rd/I during the insulin clamp) varied widely among obese participants (Figure 2A). Glucose Rd/I ranged from 181 to 382 nmol/kg FFM/min/(µU/mL) for participants in the LOW cohort and from 552 to 1274 nmol/kg FFM/min/(µU/mL) for participants in the HIGH cohort (Figure 2A and 2B). The striking differences in glucose Rd/I were present between LOW and HIGH cohorts regardless of whether glucose Rd was normalized to insulin concentration during the clamp (Figure 2C).

**Figure 2:**
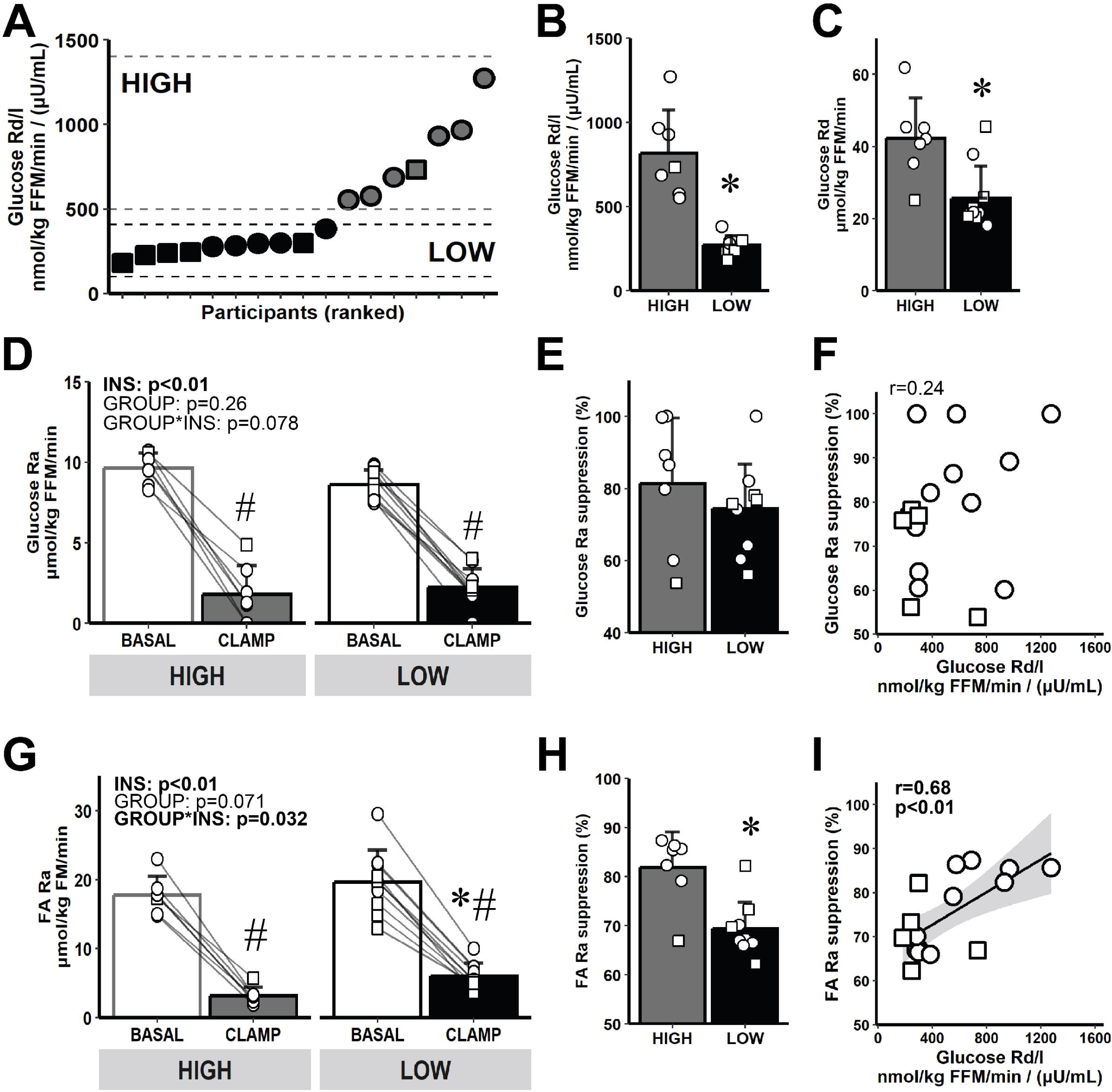
Insulin sensitivity variability across all participants and substrate kinetics. A) Glucose Rd/I variability across all participants. B-C) Stratification into HIGH (n=7) and LOW (n=10) cohorts - expressed as glucose Rd normalized to insulin [Glucose Rd/I: nmol/kg FFM/min / (µU/mL)], (B), and glucose Rd normalized only to lean body mass [(glucose Rd: µmol/kg FFM/min], (C). D) Basal and insulin-mediated glucose Ra. E) Glucose Ra % suppression. F) Correlation between glucose Rd/I and glucose Ra suppression from all participants. G) Basal and insulin-mediated FA Ra. H) FA Ra % suppression. I) Correlation between glucose Rd/I and FA Ra suppression from all participants. ○=Female, □=Male. # p<0.05 main-effect for insulin (basal versus clamp). * p<0.05 versus HIGH. Data are expressed as mean ± SD.

### Insulin effects on hepatic glucose production and FA release from adipose tissue

As expected, there was a large reduction in hepatic glucose production (glucose Ra) in response to insulin during the clamp (Figure 2D). Interestingly, despite the robust difference in insulin-mediated glucose Rd/I between LOW and HIGH, the suppression in glucose Ra (index of hepatic insulin sensitivity) was similar between groups (Figure 2E) and did not correlate with Rd/I (Figure 2F). In contrast, insulin-mediated FA Ra suppression was greater in HIGH compared with LOW (Figures 2G and 2H). Additionally, insulin-mediated FA Ra suppression was positively correlated with glucose Rd/I across all study participants (Figure 2I; r=0.68; p<0.01), suggesting a reduced ability to suppress FA mobilization from adipose tissue in response to insulin may predict an impairment in whole-body insulin-mediated glucose uptake.

### Intramyocellular LD storage in type I and II skeletal muscle fibers

Total skeletal muscle lipid content was not different between HIGH and LOW (7.8±2.0 versus 8.5±2.9 % stained, respectively; p=0.6). As anticipated, total lipid content was greater in type I versus type II muscle fibers, and this was the case for both cohorts (type I=9.3±4.9 versus type II=6.2±4.1 % stained; p<0.01; see representative image from Figure 3A). Total lipid content within the IMF and SS region of muscle fibers was also not different between HIGH and LOW (Figure 3B and 3E, respectively). There was a tendency (p=0.06) for the number of LDs per µm^2^ in the IMF region to be greater in HIGH versus LOW in both type I and II fibers (Figure 3C), but no difference was observed in the SS region (Figure 3F). LD size (measured by median LD area in each muscle fiber) within the SS region was larger in LOW versus HIGH (Figure 3G); this difference in LD area between groups was statistically significant in type II fibers (p=0.01), with a trend for a greater LD area in LOW vs HIGH in type I fibers (p=0.09). Interestingly, LD area in type II fibers was negatively correlated with insulin-mediated glucose uptake (r=-0.52, p=0.03; Figure 3H).

**Figure 3:**
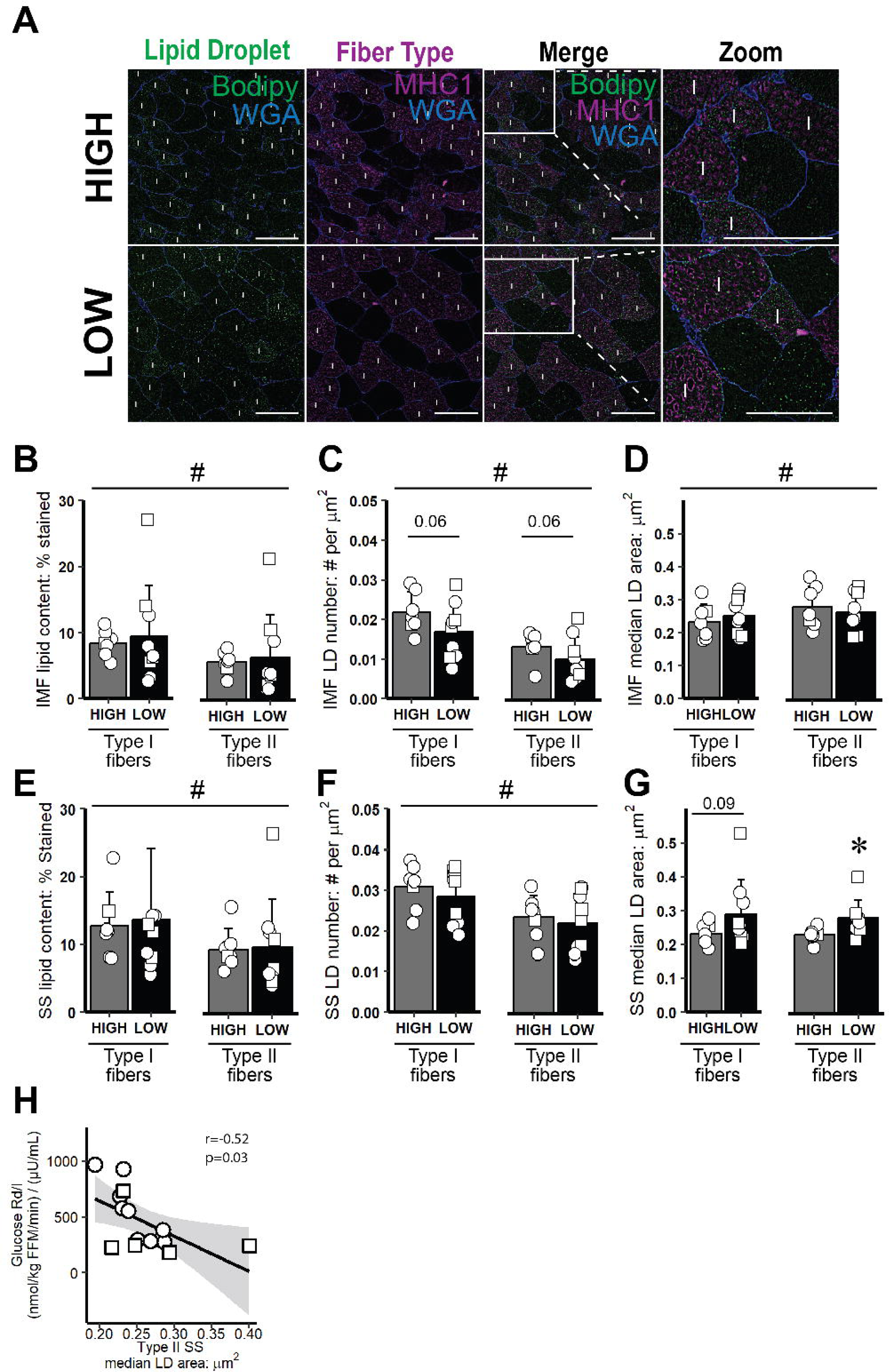
Intramyocellular LD characteristics and distribution in type I and II skeletal muscle fibers. A) Representative image in skeletal muscle of type I and II fibers from HIGH and LOW. Image contains Bodipy stain (LD, green), myofiber stain positive for MHC I (type I fibers, purple), and wheat germ agglutinin (WGA, membranes, blue). The regions inside the white boxes is enlarged for a clearer view. Magnification, x20. Scale bar, 100µm for all images. ‘I’ denotes type I muscle fiber. B-G) lipid content and LD characteristics within the IMF and SS region in type I and type II fibers. B) IMF lipid content (represented as % stained). C) IMF LD number (# LDs per µm^2^). D) IMF median LD area per fiber (µm^2^). E) SS lipid content. F) SS LD number. G) SS LD area per muscle fiber. H) Correlation between glucose Rd/I and SS LD area in type II fibers. ○=Female, □=Male; n=6 in HIGH and n=10 in LOW. # p<0.05 main effect for fiber type (type I versus type II fibers). * p<0.05 main effect for group (HIGH versus LOW), with post-hoc analysis identifying significant difference for HIGH versus LOW. Data are expressed as mean ± SD.

### Skeletal muscle insulin signaling

#### Skeletal muscle IR interaction with CD36 and Fyn

Total protein abundances for both CD36 and the Src-family kinase, Fyn, were similar between HIGH and LOW (Figures 4A and 4B), but the interaction of these proteins with the IRβ was significantly greater in HIGH versus LOW under basal and insulin-mediated conditions (Figures 4C and 4D; group main effects; p<0.01). Correlational analyses across all participants revealed that both glucose Rd/I and FA Ra suppression during the clamp were significantly correlated with the interaction of CD36 with IRβ during the clamp (both p<0.01), although correlations where not observed under basal conditions (Figures 4E-H). Glucose Rd/I during the clamp was significantly correlated with the interaction of Fyn with IRβ at basal conditions (p<0.01; Figure 4I) and during the clamp (p=0.04; Figure 4J). In contrast, insulin-mediated FA Ra suppression was not significantly correlated with the interaction of Fyn with IRβ (Figures 4K and 4L).

**Figure 4:**
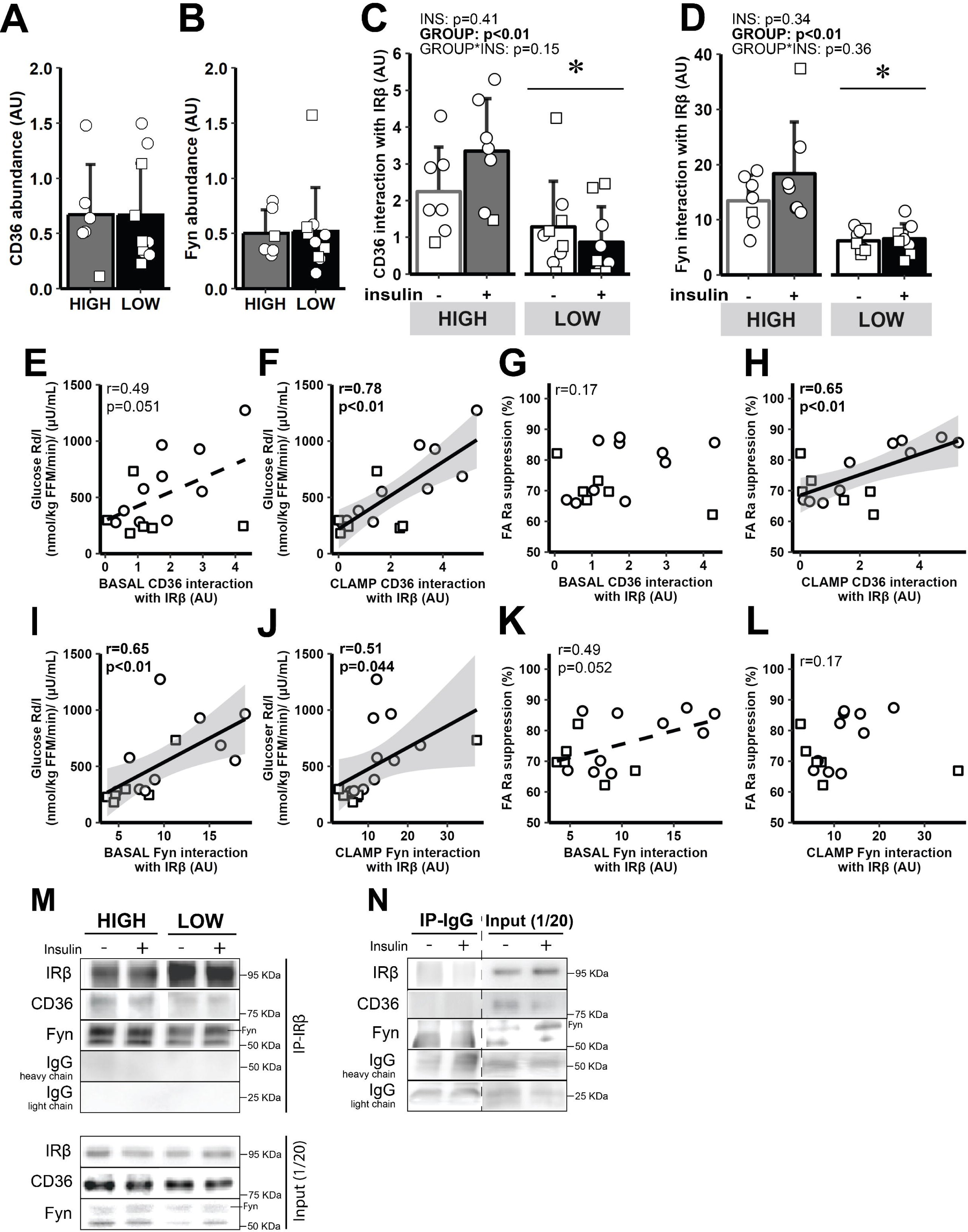
Insulin-mediated interaction between CD36 and Fyn with IRβ. A) Whole muscle CD36 abundance. B) Whole muscle Fyn abundance. C) CD36 interaction with IRβ at baseline and during the clamp. D) Fyn interaction with IRβ at baseline and during the clamp. E-F) Correlation between glucose Rd/I and CD36 interaction with IRβ at baseline (E) and during the clamp (F). G-H) Correlation between FA Ra suppression and CD36 interaction with IRβ at baseline (G) and during the clamp (H). I-J) Correlation between glucose Rd/I and Fyn interaction with IRβ at baseline (I) and during the clamp (J). K-L) Correlation between FA Ra suppression and Fyn interaction with IRβ at baseline (K) and during the clamp (L). M) Representative immunoblots from the IRβ immunoprecipitation assay, including input controls. N) IgG immunoprecipitation negative control with dashed lines to denote breaks in lanes removed for clarity. ○=Female, □=Male; HIGH versus LOW. n=7 in HIGH and n=9 in LOW. * p<0.05 - main effect for group (HIGH versus LOW). Data are expressed as mean±SD.

#### Canonical insulin signaling

Insulin-mediated phosphorylation of IRβ at Tyr^1150^ was greater in HIGH versus LOW (Figure 5A), which may align with greater Fyn tyrosine kinase interaction at IRβ. Accordingly, the insulin-mediated increase in Akt phosphorylation at Ser^473^ was also greater in HIGH versus LOW (p=0.01; Figure 5B). Insulin-stimulated Akt phosphorylation at Thr^308^ was not different between HIGH versus LOW groups (Figure 5C).

**Figure 5:**
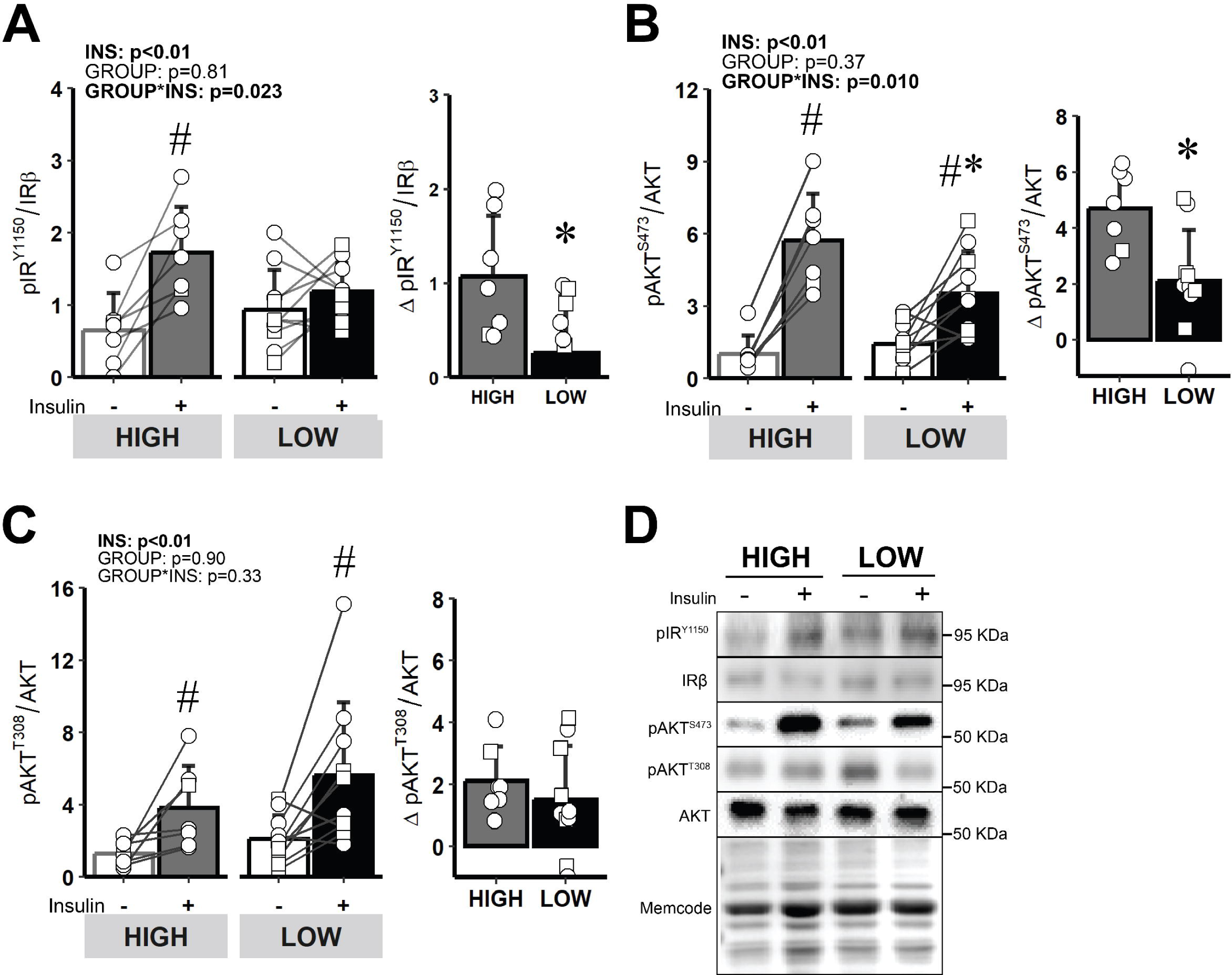
Comparison of acute insulin-mediated signaling events in skeletal muscle proximal to Akt. A) pIR^Y1150^ before and during the hyperinsulinemic clamp and Delta pIR^Y1150^ in response to insulin (difference between clamp and baseline abundance). B) pAKT^S473^ before and during the hyperinsulinemic clamp and Delta pAKT^S473^ in response to insulin. C) pAKT^T308^ before and during the hyperinsulinemic clamp and Delta pAKT^T308^ in response to insulin. D) Representative immunoblots from whole muscle lysates. ○=Female, □=Male; n=7 in HIGH and n=10 in LOW. # p<0.05 main effect for insulin (basal versus clamp). * p<0.05 versus HIGH. Data are expressed as mean ± SD.

Insulin significantly increased FOXO1 phosphorylation at Ser^256^ (p=0.01; Figure 6A) and tended to increase GSKα phosphorylation at Ser^21^ (p=0.06; Figure 6B), both of which are downstream targets of Akt. In line with the lower insulin-mediated increase in Akt phosphorylation at Ser^473^ in LOW versus HIGH, the insulin-mediated responses for phosphorylation of FOXO1 at Ser^256^ and GSKα at Ser^21^ were also blunted in LOW compared with HIGH (p<0.05; Figures 6A and 6B). Insulin did not increase AS160 phosphorylation at Thr^642^ or P44/42 MAPK (ERK 1/2) phosphorylation at Thr^202^ and Tyr^204^ in either LOW or HIGH (Figures 6C and 6D). Interestingly, we found total GLUT4 abundance to be greater in HIGH compared with LOW (p=0.01, Figure 6E).

**Figure 6:**
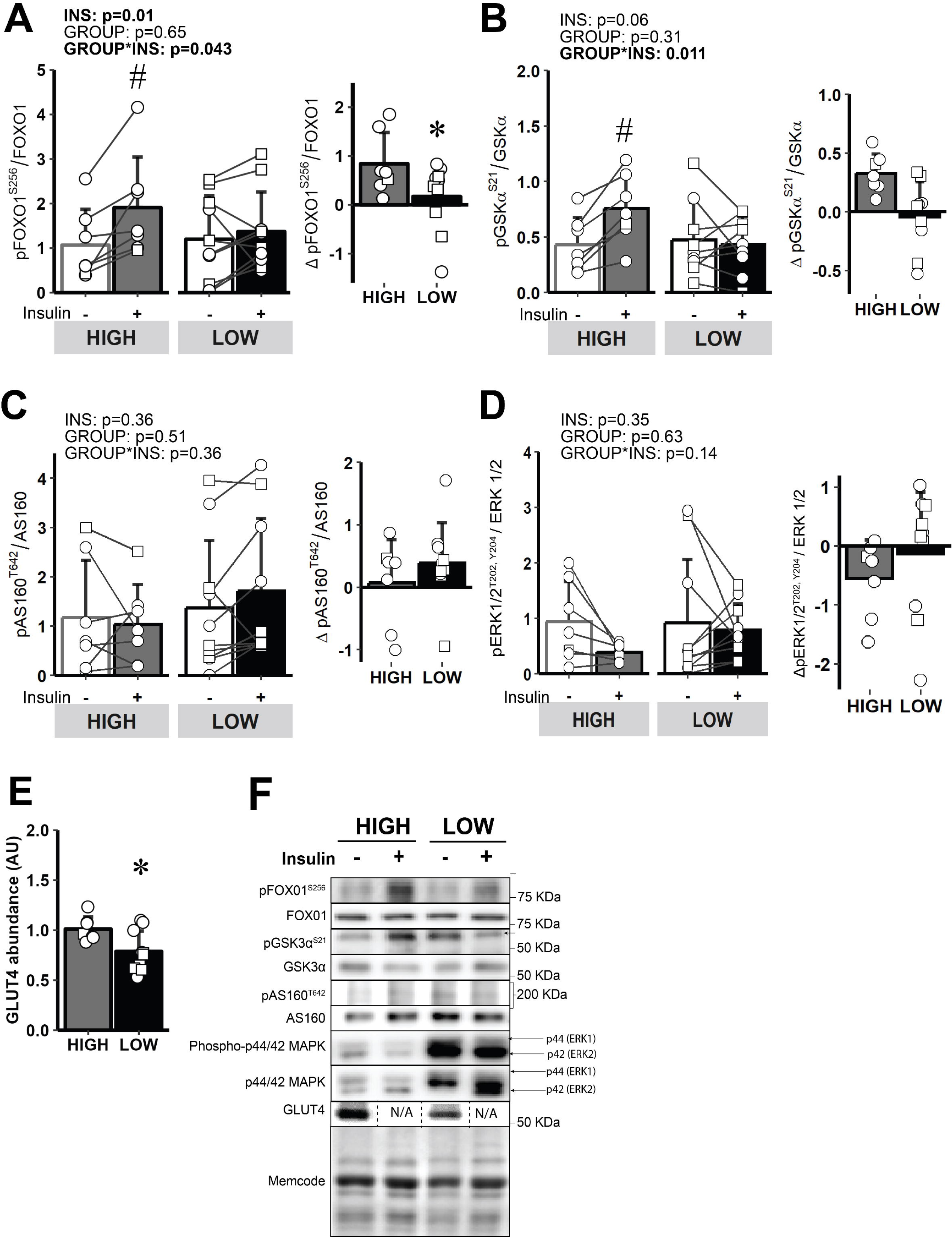
Comparison of acute insulin-mediated signaling events in skeletal muscle distal to Akt. A) pFOXO1^S256^ before and during the hyperinsulinemic clamp and Delta pFOXO1^S256^ in response to insulin. B) pGSKα^S21^ before and during the hyperinsulinemic clamp and Delta pGSKα^S21^ in response to insulin. C) pAS160^T642^ before and during the hyperinsulinemic clamp and Delta pAS160^T642^ in response to insulin. D) pERK1/2^T202/Y204^ before and during the hyperinsulinemic clamp and Delta pERK1/2^T202/Y204^ in response to insulin. E) Skeletal muscle GLUT4 abundance. F) Representative immunoblots from whole muscle. ○=Female, □=Male; n=7 in HIGH and n=10 in LOW. # p<0.05 main effect for insulin (basal versus clamp). * p<0.05 versus HIGH. Data are expressed as mean ± SD.

## DISCUSSION

Although skeletal muscle insulin resistance is very common in adults with obesity (46), not all adults with obesity are insulin resistant (4, 5), and factors underlying this difference among a relatively homogeneous cohort of adults with obesity are not well understood. Here, we confirm that a reduced ability of insulin to suppress FA mobilization from adipose tissue was a strong predictor of impaired insulin-mediated glucose uptake (7-10). This relationship is consistent with the well-described phenomenon that elevated systemic FA availability can lead to a disruption in skeletal muscle insulin signaling (9, 47-50). Studies conducted *in vitro* have reported elevated saturated FA availability reduced the interaction between CD36 and Fyn with the IR, which in turn modified insulin-mediated Tyr phosphorylation of the IR and downstream insulin signaling events (33). The interaction between CD36 and Fyn with IR was attenuated in skeletal muscle from LOW versus HIGH participants, suggesting that the impairments in skeletal muscle insulin signaling often reported in many with obesity may begin further upstream in the insulin signaling cascade than commonly considered (13, 51). Additionally, our current findings also suggest differences in the size and location of LDs within skeletal muscle may contribute to differences in insulin-mediated glucose uptake in a relatively homogeneous cohort of adults with obesity. Together, these findings support the notion that an impaired ability to suppress FA release from adipose tissue in response to insulin may lead to the development of impaired insulin signaling in skeletal muscle, perhaps in part through modifications of LD distribution within skeletal muscle, and interactions between the IR and key regulatory proteins.

The increase in plasma insulin concentration in response to a meal markedly reduces systemic FA availability in healthy adults (52, 53). In contrast, individuals who are more resistant to the effects of insulin in adipose tissue experience persistent elevations in systemic FA availability (even after meals), which is known to be a key factor underlying impaired insulin-mediated glucose uptake in skeletal muscle (9, 49, 50, 54). Because the vast majority of FA released into the systemic circulation are derived from aSAT, rather than gluteal/femoral or visceral adipose tissue (11), we contend the blunted response to insulin’s effect of reducing FA mobilization is primarily occurring in aSAT. Therefore, our findings that a blunted response to insulin’s effects on reducing FA mobilization was a primary predictor of impaired insulin-mediated peripheral glucose uptake, suggests insulin resistance in aSAT may precede the development of insulin resistance in skeletal muscle. However, the temporal pattern by which insulin resistance develops among different tissues are still not clear. Some previous reports in rodents support the notion that the onset of adipose tissue insulin resistance in response to high fat diets occurs before skeletal muscle insulin resistance (55, 56). Additionally, in lipodystrophy (a condition where adipose tissue cannot effectively store triglycerides, resulting in very high systemic FA concentrations), the excessive systemic FA availability precedes insulin resistance in skeletal muscle (57). In adults with obesity, abnormalities within aSAT include inflammation and fibrosis upon adipocyte expansion are also suggested to contribute to the development of skeletal muscle insulin resistance (58, 59).

The specific mechanisms within aSAT that may underlie an impaired ability for insulin to blunt FA mobilization are not completely understood. Insulin potently inhibits lipolytic rate, largely through Akt-mediated phosphorylation of phosphodiesterase 3B, which in turn leads to the inhibition of key lipase enzymes (60). However, it has also been proposed that low sensitivity to the anti-lipolytic response to insulin may not be responsible for the blunted FA Ra suppression in response to insulin (61). Because adipose tissue is often exceptionally sensitive to the anti-lipolytic effects of insulin, it is possible that a modest impairment of insulin signaling in aSAT may not lead to a detectable difference in lipolytic rate. Alternatively, a modest impairment in insulin signaling in adipocytes can impact other processes, such as the rate of FA re-esterification (62, 63), which may be an important contributor to the blunted FA Ra suppression reported here. Local inflammation (e.g., TNFα, IL6, and IL1β) within adipose tissue has been attributed to impair insulin signaling in adipose tissue (64-66), and may contribute to impaired ability to reduce FA mobilization in response to insulin. Morphological features of aSAT such as large adipocytes may also contribute to a blunted suppression in FA release, perhaps due in part to greater pro-inflammatory immune cell infiltration (67). In addition, relatively low capillary density coupled with hypertrophied adipocytes can result in attenuated insulin delivery, as well as induce adipocyte hypoxia, which has been linked to impaired insulin responsiveness (58, 59). Excessive fibrosis within the ECM of aSAT has also been linked with an impaired response to insulin in adipose tissue (68, 69). How aSAT fibrotic content or composition may reduce insulin signaling within adipose tissue remains elusive, but potential factors include increased expression/production of pro-inflammatory cytokines by mechanical stress induced by excess collagen accumulation (70), and physical constraints limiting insulin delivery to adipocytes (68, 71). Overall, excessive fibrosis, large adipocyte size, relatively low capillary density, greater inflammatory profile, and attenuated insulin signaling in aSAT may converge to blunt insulin’s ability to suppress FA mobilization, and in turn, a resultant chronic elevation in systemic FA may lead to impaired insulin-mediated glucose uptake in skeletal muscle.

Insulin resistance in skeletal muscle stemming from high FA availability has been causally linked to DAG and ceramide accumulation (17, 20), resulting in non-typical PKC isoform activation (72), and decreased IRS-1 and AKT activation (13, 51, 73). However, there remains ongoing debate regarding lipid-mediated disruption of canonical insulin signaling in skeletal muscle (23-25). The membrane glycoprotein CD36 is well-known for its role in long-chain FA transport, but recent evidence demonstrated CD36 can also modify insulin signaling (33, 38). The physical interaction between CD36 and the IR was found to enhance IR phosphorylation by recruiting the Src-family kinase protein, Fyn, to the IR where activation via Tyr phosphorylation occurs (33). Our findings that the interaction between the IR and CD36 and Fyn was greater in HIGH versus LOW align with the notion that a greater interaction among these proteins may help propagate intracellular insulin action. Importantly, elevated saturated FA availability was previously found to attenuate CD36 from recruiting Fyn to the IR, contributing to the blunted insulin-mediated glucose uptake commonly found when FA availability is high (33). Therefore, the reduced ability of insulin to suppress FA mobilization (leading to elevated FA availability) in our LOW participants may have blunted the ability of CD36 to recruit Fyn to the IR, reducing the interaction between Fyn and the IR, leading to an attenuation in IR phosphorylation. Interestingly, the elevation in plasma insulin concentration during the clamp in our study did not significantly increase the interaction of either CD36 or Fyn with the IR, suggesting that the basal (i.e., non-insulin stimulated) interactions of these proteins may be important in the regulation of subsequent IR phosphorylation in response to insulin. Our findings that insulin-mediated phosphorylation of AKT at Ser^473^, as well as phosphorylation of the Akt substrates, GSK3α at Ser^21^ and FOXO1 at Ser^256^ were lower in LOW versus HIGH, align with the notion that interaction between the IR and Fyn may impact canonical insulin signaling downstream of the IR. Whether the CD36-mediated recruitment of Fyn to the IR involves a direct physical interaction between CD36 and Fyn is not known. However, Fyn binding sites have been identified along the cytoplasmic lipid raft domains of CD36 (74, 75), suggesting this process might involve direct binding between these proteins for CD36 to allow phosphorylation of the IR by Fyn. Unfortunately, the relatively large amount of tissue required for the co-immunoprecipitation assay limited our ability to confirm whether CD36 interaction with Fyn was also attenuated in LOW versus HIGH. Together, these findings are the first to our knowledge in human skeletal muscle to support a previously proposed mechanism by which CD36 regulates IR phosphorylation (33) and downstream signaling (38), and that FA availability and uptake into skeletal muscle may modify this effect.

Lipid accumulation in skeletal muscle LDs, and LD location within the myocyte have also been implicated in the impairment of insulin-mediated glucose uptake in skeletal muscle (76). Our observation that LDs in the SS region of the myocyte were larger in LOW versus HIGH, aligns with previous reports demonstrating correlations between SS LD size and insulin resistance (30-32, 77). The mechanistic link explaining how the over-accumulation of larger LDs in the SS region of the myocyte induces insulin resistance is still unclear, but the proximity of these large LDs to the muscle membrane may disrupt key insulin signaling processes (30). For example, the accumulation of lipid intermediates in the SS region, which may be released from SS LDs, have been linked with insulin resistance (16, 78). Additionally, larger LDs have a lower surface area-to-volume ratios than small LDs, which may limit lipolytic control at the membrane compared to smaller LDs. In turn, this may increase the susceptibility for incomplete lipolysis (79-81), resulting in aberrant lipid accumulation within the SS region of the myocyte, in close proximity to insulin signaling events that occur at/near the plasma membrane. Our findings that the number of LDs in the IMF region tended to be greater in HIGH versus LOW suggests that a partitioning of LDs away from the SS region may be protective against impaired insulin signaling. The greater abundance of LDs in the IMF region may also help protect against the development of insulin resistance by their close proximity to more mitochondria, which may aid in the direct lipid transfer towards oxidative metabolism, and thereby may mitigate cytosolic lipid accumulation (29, 82). We contend that differences in LD size and distribution observed between LOW versus HIGH may be due in part to differences in the ability of insulin to suppress FA release from adipose tissue, whereby, a chronic elevation in systemic FA availability and uptake in individuals with low FA Ra suppression in response to insulin modifies LD storage and metabolism.

Although we interpret our findings describing the relationship between insulin-mediated suppression of FA release and glucose uptake to suggest a suppressed response to insulin in adipose tissue may contribute the development of whole-body insulin resistance, we acknowledge that this interpretation was largely based on correlational analyses that do not direct causation. However, these findings are consistent with other studies supporting the observation that insulin resistance development is likely a consequence of excess FA release from aSAT (7-10), likely resulting in greater FA uptake into skeletal muscle. We also acknowledge that our *in vivo* measurements aimed at identifying key contributors to insulin resistance in our participants were not exhaustive, and additional factors may contribute to insulin resistance (e.g., vascular dysfunction limiting insulin delivery) (83, 84). Additionally, the phosphorylation of canonical insulin signaling can also be regulated by other intracellular responses that were not measured in this study (e.g., PKCs, JNK, and mTORC1/S6 kinase) – many of which are known to negatively influence activation of proximal insulin signaling components (85). It is also important to acknowledge that our LOW group had a greater proportion of male participants compared to HIGH (LOW: 5M/5F, HIGH: 6F/1M) and men are commonly found to be more insulin resistant than women (86, 87). Therefore, we cannot exclude the possibility that sex differences are contributing to differences observed in FA release and whole-body insulin sensitivity between groups.

In conclusion, our findings suggest that a reduced ability to suppress FA mobilization from aSAT in response to insulin is an important contributor to whole-body insulin resistance. A blunted insulin-induced suppression of FA mobilization can lead to a persistent elevation in systemic FA availability, which is a major factor underlying the development of insulin resistance in obesity. Participants in our study with relatively low insulin-mediated glucose uptake also presented larger-sized LDs in the SS region of the myocyte, which may contribute to their insulin resistance by interfering with key insulin signaling processes at the muscle membrane. Novel findings from our study also suggest this high systemic FA availability may diminish skeletal muscle insulin signaling, in part by reducing the interaction between CD36 and Fyn with the IR, thereby lowering downstream insulin signaling. Overall, our findings suggest that attenuated suppression in FA mobilization by insulin may contribute to a tissue-specific crosstalk, whereby elevated FA release from aSAT modifies skeletal muscle LD size and distribution in the myocyte and disrupts IR interaction with key regulatory proteins, leading to skeletal muscle insulin resistance.

## REFERENCES

1. Hales CM, Carroll MD, Fryar CD, and Ogden CL. Prevalence of Obesity and Severe Obesity Among Adults: United States, 2017-2018. NCHS Data Brief. 2020(360):1-8.

2. Farin HM, Abbasi F, and Reaven GM. Body mass index and waist circumference both contribute to differences in insulin-mediated glucose disposal in nondiabetic adults. Am J Clin Nutr. 2006;83(1):47–51.

3. Reaven GM. Banting lecture 1988. Role of insulin resistance in human disease. Diabetes. 1988;37(12):1595-607.

4. Smith GI, Mittendorfer B, and Klein S. Metabolically healthy obesity: facts and fantasies. J Clin Invest. 2019;129(10):3978–89.

5. Wildman RP, Muntner P, Reynolds K, McGinn AP, Rajpathak S, Wylie-Rosett J, et al. The Obese Without Cardiometabolic Risk Factor Clustering and the Normal Weight With Cardiometabolic Risk Factor Clustering: Prevalence and Correlates of 2 Phenotypes Among the US Population (NHANES 1999-2004). Arch Intern Med. 2008;168(15):1617–24.

6. Van Pelt DW, Newsom SA, Schenk S, and Horowitz JF. Relatively low endogenous fatty acid mobilization and uptake helps preserve insulin sensitivity in obese women. Int J Obes (Lond). 2015;39(1):149–55.

7. Schleh MW, Ryan BJ, Ahn C, Ludzki AC, Varshney P, Gillen JB, et al. Metabolic dysfunction in obesity is related to impaired suppression of fatty acid release from adipose tissue by insulin. Obesity (Silver Spring, Md). 2023;31(5):1347–61.

8. Magkos F, Fabbrini E, Conte C, Patterson BW, and Klein S. Relationship between Adipose Tissue Lipolytic Activity and Skeletal Muscle Insulin Resistance in Nondiabetic Women. J Clin Endocrinol Metab. 2012;97(7):E1219–E23.

9. Groop LC, Bonadonna RC, DelPrato S, Ratheiser K, Zyck K, Ferrannini E, et al. Glucose and free fatty acid metabolism in non-insulin-dependent diabetes mellitus. Evidence for multiple sites of insulin resistance. J Clin Invest. 1989;84(1):205–13.

10. Koh H-CE, van Vliet S, Pietka TA, Meyer GA, Razani B, Laforest R, et al. Subcutaneous adipose tissue metabolic function and insulin sensitivity in people with obesity. Diabetes. 2021:db210160.

11. Nielsen S, Guo Z, Johnson CM, Hensrud DD, and Jensen MD. Splanchnic lipolysis in human obesity. J Clin Invest. 2004;113(11):1582–8.

12. Krssak M, Falk Petersen K, Dresner A, DiPietro L, Vogel SM, Rothman DL, et al. Intramyocellular lipid concentrations are correlated with insulin sensitivity in humans: a 1H NMR spectroscopy study. Diabetologia. 1999;42(1):113–6.

13. Dresner A, Laurent D, Marcucci M, Griffin ME, Dufour S, Cline GW, et al. Effects of free fatty acids on glucose transport and IRS-1-associated phosphatidylinositol 3-kinase activity. The Journal of clinical investigation. 1999;103(2):253–9.

14. Weiss R, Dufour S, Taksali SE, Tamborlane WV, Petersen KF, Bonadonna RC, et al. Prediabetes in obese youth: a syndrome of impaired glucose tolerance, severe insulin resistance, and altered myocellular and abdominal fat partitioning. The Lancet. 2003;362(9388):951-7.

15. DeVito LM, Dennis EA, Kahn BB, Shulman GI, Witztum JL, Sadhu S, et al. Bioactive lipids and metabolic syndrome—a symposium report. Annals of the New York Academy of Sciences. 2022;1511(1):87–106.

16. Song JD, Alves TC, Befroy DE, Perry RJ, Mason GF, Zhang X-M, et al. Dissociation of Muscle Insulin Resistance from Alterations in Mitochondrial Substrate Preference. Cell Metabolism. 2020;32(5):726–35.e5.

17. Adams JM, Pratipanawatr T, Berria R, Wang E, DeFronzo RA, Sullards MC, et al. Ceramide Content Is Increased in Skeletal Muscle From Obese Insulin-Resistant Humans. Diabetes. 2004;53(1):25–31.

18. Koves TR, Ussher JR, Noland RC, Slentz D, Mosedale M, Ilkayeva O, et al. Mitochondrial Overload and Incomplete Fatty Acid Oxidation Contribute to Skeletal Muscle Insulin Resistance. Cell Metab. 2008;7(1):45–56.

19. Houmard JA, Tanner CJ, Yu C, Cunningham PG, Pories WJ, MacDonald KG, et al. Effect of weight loss on insulin sensitivity and intramuscular long-chain fatty acyl-CoAs in morbidly obese subjects. Diabetes. 2002;51(10):2959–63.

20. Itani SI, Ruderman NB, Schmieder F, and Boden G. Lipid-Induced Insulin Resistance in Human Muscle Is Associated With Changes in Diacylglycerol, Protein Kinase C, and IκB-α. Diabetes. 2002;51(7):2005–11.

21. Batista TM, Haider N, and Kahn CR. Defining the underlying defect in insulin action in type 2 diabetes. Diabetologia. 2021;64(5):994–1006.

22. Moeschel K, Beck A, Weigert C, Lammers R, Kalbacher H, Voelter W, et al. Protein kinase C-zeta-induced phosphorylation of Ser318 in insulin receptor substrate-1 (IRS-1) attenuates the interaction with the insulin receptor and the tyrosine phosphorylation of IRS-1. J Biol Chem. 2004;279(24):25157–63.

23. Hoy AJ, Brandon AE, Turner N, Watt MJ, Bruce CR, Cooney GJ, et al. Lipid and insulin infusion-induced skeletal muscle insulin resistance is likely due to metabolic feedback and not changes in IRS-1, Akt, or AS160 phosphorylation. American Journal of Physiology-Endocrinology and Metabolism. 2009;297(1):E67-E75.

24. Skovbro M, Baranowski M, Skov-Jensen C, Flint A, Dela F, Gorski J, et al. Human skeletal muscle ceramide content is not a major factor in muscle insulin sensitivity. Diabetologia. 2008;51(7):1253–60.

25. Timmers S, Nabben M, Bosma M, van Bree B, Lenaers E, van Beurden D, et al. Augmenting muscle diacylglycerol and triacylglycerol content by blocking fatty acid oxidation does not impede insulin sensitivity. Proceedings of the National Academy of Sciences of the United States of America. 2012;109(29):11711–6.

26. Walther TC, and Farese RV, Jr. Lipid droplets and cellular lipid metabolism. Annu Rev Biochem. 2012;81:687–714.

27. Olzmann JA, and Carvalho P. Dynamics and functions of lipid droplets. Nat Rev Mol Cell Biol. 2019;20(3):137–55.

28. Ferreira R, Vitorino R, Alves RM, Appell HJ, Powers SK, Duarte JA, et al. Subsarcolemmal and intermyofibrillar mitochondria proteome differences disclose functional specializations in skeletal muscle. Proteomics. 2010;10(17):3142–54.

29. Shaw CS, Jones DA, and Wagenmakers AJM. Network distribution of mitochondria and lipid droplets in human muscle fibres. Histochem Cell Biol. 2008;129(1):65–72.

30. Daemen S, Gemmink A, Brouwers B, Meex RCR, Huntjens PR, Schaart G, et al. Distinct lipid droplet characteristics and distribution unmask the apparent contradiction of the athlete’s paradox. Mol Metab. 2018;17:71–81.

31. Nielsen J, Christensen AE, Nellemann B, and Christensen B. Lipid droplet size and location in human skeletal muscle fibers are associated with insulin sensitivity. Am J Physiol Endocrinol Metab. 2017;313(6):E721–E30.

32. Nielsen J, Mogensen M, Vind BF, Sahlin K, Højlund K, Schrøder HD, et al. Increased subsarcolemmal lipids in type 2 diabetes: effect of training on localization of lipids, mitochondria, and glycogen in sedentary human skeletal muscle. Am J Physiol Endocrinol Metab. 2010;298(3):E706–13.

33. Samovski D, Dhule P, Pietka T, Jacome-Sosa M, Penrose E, Son N-H, et al. Regulation of Insulin Receptor Pathway and Glucose Metabolism by CD36 Signaling. 2018;67(7):1272-84.

34. Pepino MY, Kuda O, Samovski D, and Abumrad NA. Structure-function of CD36 and importance of fatty acid signal transduction in fat metabolism. Annu Rev Nutr. 2014;34:281–303.

35. Su X, and Abumrad NA. Cellular fatty acid uptake: a pathway under construction. Trends Endocrinol Metab. 2009;20(2):72–7.

36. Samovski D, Jacome-Sosa M, and Abumrad NA. Fatty Acid Transport and Signaling: Mechanisms and Physiological Implications. Annual Review of Physiology. 2023;85(1):317–37.

37. Glatz JC, and Luiken JFP. Dynamic role of the transmembrane glycoprotein CD36 (SR-B2) in cellular fatty acid uptake and utilization. Journal of Lipid Research. 2018;59(7):1084–93.

38. Sun S, Tan P, Huang X, Zhang W, Kong C, Ren F, et al. Ubiquitinated CD36 sustains insulin-stimulated Akt activation by stabilizing insulin receptor substrate 1 in myotubes. Journal of Biological Chemistry. 2018;293(7):2383–94.

39. Samovski D, Sun J, Pietka T, Gross RW, Eckel RH, Su X, et al. Regulation of AMPK Activation by CD36 Links Fatty Acid Uptake to β-Oxidation. 2015;64(2):353-9.

40. DeFronzo RA, Tobin JD, and Andres R. Glucose clamp technique: a method for quantifying insulin secretion and resistance. Am J Physiol. 1979;237(3):E214–23.

41. Ascaso JF, Pardo S, Real JT, Lorente RI, Priego A, and Carmena R. Diagnosing Insulin Resistance by Simple Quantitative Methods in Subjects With Normal Glucose Metabolism. Diabetes Care. 2003;26(12):3320–5.

42. Reeder SB, Cruite I, Hamilton G, and Sirlin CB. Quantitative Assessment of Liver Fat with Magnetic Resonance Imaging and Spectroscopy. J Magn Reson Imaging. 2011;34(4):729–49.

43. Ryan BJ, Schleh MW, Ahn C, Ludzki AC, Gillen JB, Varshney P, et al. Moderate-intensity exercise and high-intensity interval training affect insulin sensitivity similarly in obese adults. J Clin Endocrinol Metab. 2020.

44. Newsom SA, Schenk S, Thomas KM, Harber MP, Knuth ND, Goldenberg N, et al. Energy deficit after exercise augments lipid mobilization but does not contribute to the exercise-induced increase in insulin sensitivity. 2010;108(3):554–60.

45. Steele R. Influences of glucose loading and of injected insulin on hepatic glucose output. Ann N Y Acad Sci. 1959;82:420–30.

46. DeFronzo RA, and Ferrannini E. Insulin Resistance: A Multifaceted Syndrome Responsible for NIDDM, Obesity, Hypertension, Dyslipidemia, and Atherosclerotic Cardiovascular Disease. Diabetes Care. 1991;14(3):173–94.

47. Roden M, Price TB, Perseghin G, Petersen KF, Rothman DL, Cline GW, et al. Mechanism of free fatty acid-induced insulin resistance in humans. The Journal of clinical investigation. 1996;97(12):2859–65.

48. Roden M, Stingl H, Chandramouli V, Schumann WC, Hofer A, Landau BR, et al. Effects of free fatty acid elevation on postabsorptive endogenous glucose production and gluconeogenesis in humans. Diabetes. 2000;49(5):701–7.

49. Boden G. Role of fatty acids in the pathogenesis of insulin resistance and NIDDM. Diabetes. 1997;46(1):3–10.

50. Thiébaud D, DeFronzo RA, Jacot E, Golay A, Acheson K, Maeder E, et al. Effect of long chain triglyceride infusion on glucose metabolism in man. Metabolism. 1982;31(11):1128–36.

51. Li Y, Soos TJ, Li X, Wu J, DeGennaro M, Sun X, et al. Protein Kinase C θ Inhibits Insulin Signaling by Phosphorylating IRS1 at Ser1101. Journal of Biological Chemistry. 2004;279(44):45304–7.

52. Coppack SW, Fisher RM, Gibbons GF, Humphreys SM, McDonough MJ, Potts JL, et al. Postprandial substrate deposition in human forearm and adipose tissues in vivo. Clin Sci (Lond). 1990;79(4):339–48.

53. Frayn KN. Adipose tissue as a buffer for daily lipid flux. Diabetologia. 2002;45(9):1201–10.

54. Roden M, Krssak M, Stingl H, Gruber S, Hofer A, Fürnsinn C, et al. Rapid impairment of skeletal muscle glucose transport/phosphorylation by free fatty acids in humans. Diabetes. 1999;48(2):358–64.

55. Turner N, Kowalski GM, Leslie SJ, Risis S, Yang C, Lee-Young RS, et al. Distinct patterns of tissue-specific lipid accumulation during the induction of insulin resistance in mice by high-fat feeding. Diabetologia. 2013;56(7):1638–48.

56. Burchfield JG, Kebede MA, Meoli CC, Stöckli J, Whitworth PT, Wright AL, et al. High dietary fat and sucrose result in an extensive and time-dependent deterioration in health of multiple physiological systems in mice. Journal of Biological Chemistry. 2018;293(15):5731–45.

57. Mann JP, and Savage DB. What lipodystrophies teach us about the metabolic syndrome. The Journal of Clinical Investigation. 2019;129(10):4009–21.

58. Cifarelli V, Beeman SC, Smith GI, Yoshino J, Morozov D, Beals JW, et al. Decreased adipose tissue oxygenation associates with insulin resistance in individuals with obesity. J Clin Invest. 2020;130(12):6688–99.

59. Crewe C, An YA, and Scherer PE. The ominous triad of adipose tissue dysfunction: inflammation, fibrosis, and impaired angiogenesis. J Clin Invest. 2017;127(1):74–82.

60. DiPilato LM, Ahmad F, Harms M, Seale P, Manganiello V, and Birnbaum MJ. The Role of PDE3B Phosphorylation in the Inhibition of Lipolysis by Insulin. Mol Cell Biol. 2015;35(16):2752–60.

61. Yeckel CW, Dziura J, and DiPietro L. Abdominal obesity in older women: potential role for disrupted fatty acid reesterification in insulin resistance. J Clin Endocrinol Metab. 2008;93(4):1285–91.

62. Meegalla RL, Billheimer JT, and Cheng D. Concerted elevation of acyl-coenzyme A:diacylglycerol acyltransferase (DGAT) activity through independent stimulation of mRNA expression of DGAT1 and DGAT2 by carbohydrate and insulin. Biochem Biophys Res Commun. 2002;298(3):317–23.

63. Ali AH, Mundi M, Koutsari C, Bernlohr DA, and Jensen MD. Adipose Tissue Free Fatty Acid Storage In Vivo: Effects of Insulin Versus Niacin as a Control for Suppression of Lipolysis. Diabetes. 2015;64(8):2828–35.

64. Lagathu C, Yvan-Charvet L, Bastard JP, Maachi M, Quignard-Boulangé A, Capeau J, et al. Long-term treatment with interleukin-1β induces insulin resistance in murine and human adipocytes. Diabetologia. 2006;49(9):2162–73.

65. Hotamisligil GS, Peraldi P, Budavari A, Ellis R, White MF, and Spiegelman BM. IRS-1-mediated inhibition of insulin receptor tyrosine kinase activity in TNF-alpha- and obesity-induced insulin resistance. Science. 1996;271(5249):665-8.

66. Lagathu C, Bastard J-P, Auclair M, Maachi M, Capeau J, and Caron M. Chronic interleukin-6 (IL-6) treatment increased IL-6 secretion and induced insulin resistance in adipocyte: prevention by rosiglitazone. Biochem Biophys Res Commun. 2003;311(2):372–9.

67. Guilherme A, Virbasius JV, Puri V, and Czech MP. Adipocyte dysfunctions linking obesity to insulin resistance and type 2 diabetes. Nat Rev Mol Cell Biol. 2008;9(5):367–77.

68. Marcelin G, Ferreira A, Liu Y, Atlan M, Aron-Wisnewsky J, Pelloux V, et al. A PDGFRα-Mediated Switch toward CD9(high) Adipocyte Progenitors Controls Obesity-Induced Adipose Tissue Fibrosis. Cell Metab. 2017;25(3):673–85.

69. Khan T, Muise ES, Iyengar P, Wang ZV, Chandalia M, Abate N, et al. Metabolic Dysregulation and Adipose Tissue Fibrosis: Role of Collagen VI. Mol Cell Biol. 2009;29(6):1575–91.

70. Pellegrinelli V, Heuvingh J, du Roure O, Rouault C, Devulder A, Klein C, et al. Human adipocyte function is impacted by mechanical cues. J Pathol. 2014;233(2):183–95.

71. Marcelin G, Silveira ALM, Martins LB, Ferreira AVM, and Clément K. Deciphering the cellular interplays underlying obesity-induced adipose tissue fibrosis. J Clin Invest. 2019;129(10):4032–40.

72. Itani SI, Zhou Q, Pories WJ, MacDonald KG, and Dohm GL. Involvement of protein kinase C in human skeletal muscle insulin resistance and obesity. Diabetes. 2000;49(8):1353–8.

73. Dobrowsky RT, Kamibayashi C, Mumby MC, and Hannun YA. Ceramide activates heterotrimeric protein phosphatase 2A. J Biol Chem. 1993;268(21):15523–30.

74. Huang MM, Bolen JB, Barnwell JW, Shattil SJ, and Brugge JS. Membrane glycoprotein IV (CD36) is physically associated with the Fyn, Lyn, and Yes protein-tyrosine kinases in human platelets. Proceedings of the National Academy of Sciences of the United States of America. 1991;88(17):7844–8.

75. Bull HA, Brickell PM, and Dowd PM. Src-related protein tyrosine kinases are physically associated with the surface antigen CD36 in human dermal microvascular endothelial cells. FEBS letters. 1994;351(1):41–4.

76. Gemmink A, Goodpaster BH, Schrauwen P, and Hesselink MKC. Intramyocellular lipid droplets and insulin sensitivity, the human perspective. Biochimica et Biophysica Acta (BBA) - Molecular and Cell Biology of Lipids. 2017;1862(10, Part B):1242-9.

77. Chee C, Shannon CE, Burns A, Selby AL, Wilkinson D, Smith K, et al. Relative Contribution of Intramyocellular Lipid to Whole-Body Fat Oxidation Is Reduced With Age but Subsarcolemmal Lipid Accumulation and Insulin Resistance Are Only Associated With Overweight Individuals. Diabetes. 2016;65(4):840–50.

78. Perreault L, Newsom SA, Strauss A, Kerege A, Kahn DE, Harrison KA, et al. Intracellular localization of diacylglycerols and sphingolipids influences insulin sensitivity and mitochondrial function in human skeletal muscle. JCI Insight. 2018;3(3):e96805.

79. Thiam AR, and Beller M. The why, when and how of lipid droplet diversity. J Cell Sci. 2017;130(2):315–24.

80. Murphy DJ. The dynamic roles of intracellular lipid droplets: from archaea to mammals. Protoplasma. 2012;249(3):541–85.

81. Hesselink MK, Mensink M, and Schrauwen P. Intramyocellular lipids and insulin sensitivity: does size really matter? Obes Res. 2004;12(5):741–2.

82. Vock R, Hoppeler H, Claassen H, Wu DX, Billeter R, Weber JM, et al. Design of the oxygen and substrate pathways. VI. structural basis of intracellular substrate supply to mitochondria in muscle cells. J Exp Biol. 1996;199(Pt 8):1689–97.

83. Clerk LH, Vincent MA, Jahn LA, Liu Z, Lindner JR, and Barrett EJ. Obesity blunts insulin-mediated microvascular recruitment in human forearm muscle. Diabetes. 2006;55(5):1436–42.

84. McClatchey PM, Williams IM, Xu Z, Mignemi NA, Hughey CC, McGuinness OP, et al. Perfusion controls muscle glucose uptake by altering the rate of glucose dispersion in vivo. Am J Physiol Endocrinol Metab. 2019;317(6):E1022–e36.

85. Fazakerley DJ, Krycer JR, Kearney AL, Hocking SL, and James DE. Muscle and adipose tissue insulin resistance: malady without mechanism? Journal of Lipid Research. 2019;60(10):1720–32.

86. Nuutila P, Knuuti MJ, Mäki M, Laine H, Ruotsalainen U, Teräs M, et al. Gender and insulin sensitivity in the heart and in skeletal muscles. Studies using positron emission tomography. Diabetes. 1995;44(1):31–6.

87. Tramunt B, Smati S, Grandgeorge N, Lenfant F, Arnal J-F, Montagner A, et al. Sex differences in metabolic regulation and diabetes susceptibility. Diabetologia. 2020;63(3):453–61.

