## Supplementary material for "Insulin-mediated suppression of fatty acid release predicts whole-body insulin resistance of glucose uptake and skeletal muscle insulin receptor activation": Table S1

### SUPPLEMENTARY INFORMATION

| Table S1: Antibodies and Reagents | | |  |
| --- | --- | --- | --- |
| **Primary and secondary antibodies** | **Source** | **Identifier** | **Concentration** |
| Phospho-IGF-I Receptor β (Tyr1135/1136)/Insulin Receptor β (Tyr1150/1151) | CST | 3024 | 1:1000 |
| anti-Insulin Receptor β | CST | 3025 | 1:1000 |
| anti-pAkt^S473^ | CST | 9271 | 1:1000 |
| anti-pAkt^T308^ | CST | 9275 | 1:1000 |
| anti-Akt | CST | 9272 | 1:2000 |
| pGSK3α^S21^ | CST | 4337 | 1:1000 |
| anti-GSK3α | CST | 5676 | 1:1000 |
| anti-pFOXO1^S256^ | CST | 9461 | 1:1000 |
| anti-FoxO1 | CST | 9454 | 1:1000 |
| anti-pAS160^T642^ | CST | 8881 | 1:500 |
| anti-AS160 | CST | 2670 | 1:1000 |
| anti-human CD36 | R&D Systems | AF1955 | 1:500 |
| anti-Fyn | CST | 4023 | 1:1000 |
| anti-GLUT4 | Santa Cruz | sc-53566 | 1:1000 |
| Phospho-p44^Thr202^/42^Tyr204^ MAPK (Erk1/2) | CST | 9101 | 1:1000 |
| p44/42 MAPK (Erk1/2) | CST | 9102 | 1:1000 |
| Normal Rabbit IgG | CST | 2729 | 1:1000 |
| anti-rabbit IgG, HRP-linked Antibody | CST | 7074 | 1:10000 |
| anti-mouse IgG, HRP-linked Antibody | CST | 7076 | 1:10000 |
| **Stable Isotope Tracers** |  |  |  |
| D-Glucose (6,6-D2) | Cambridge Isotopes | DLM-349-PK | N/A |
| Potassium Palmitate (1-13C, 99%) | Cambridge Isotopes | CLM-1889-PK | N/A |
| **Histochemistry Reagents** |  |  |  |
| BODIPY 493/503 | Invitrogen Thermo-Fisher | D3922 | 1:100 |
| BA-D5: Myosin heavy chain Type 1 | Developmental Studies Hydroma Bank | - | 1:200 |
| Alexa Fluor 647 Goat anti-Mouse IgG2b | Invitrogen Thermo-Fisher | A21242 | 1:200 |
| Wheat Germ Agglutinin - Alexa Fluor 555 | Invitrogen Thermo-Fisher | W32464 | 1:500 |
| Triton X-100 | Sigma-Aldrich | T8787 | N/A |
| ProLong Gold Antifade Mountant | Invitrogen Thermo-Fisher | P36930 | N’A |
| **Plasma Analysis Reagents** |  |  |  |
| Glucose Oxidase | Thermo-Fisher | A22189 | N/A |
| NEFA Standard Solution | Wako Diagnostics | 27676491 | N/A |
| Triglyceride Reagent | Sigma-Aldrich | T2449 | N/A |
| HDL-Cholesterol E | Wako Diagnostics | 9059676 | N/A |
| Total Cholesterol E | Wako Diagnostics | 9138103 | N/A |
| Human IL-6 ELISA | R&D Systems | HS600C | N/A |
| C-Reactive Protein ELISA | Calbiotech | CR120C | N/A |
| Human Leptin ELISA | Millipore Sigma | EZHL80SK | N/A |
| Human Total Adiponectin/Acrp30 ELISA | R&D Systems | DRP300 | N/A |
| Human HMW Adiponectin/Acrp30 ELISA | R&D Systems | DHWAD0 | N/A |
| Insulin For Immulite | Siemens | 10381429 | N/A |

Abbreviations: CST, Cell Signaling Technology
